# Discovery of new deregulated miRNAs in gingivo buccal carcinoma using Group Benjamini Hochberg method: a commentary on “A quest for miRNA bio-marker: a track back approach from gingivo buccal cancer to two different types of precancers”

**DOI:** 10.1101/2023.02.17.529013

**Authors:** Salil Koner, Navonil De Sarkar, Nilanjana Laha

**Author notes:** Senior author (Nilanjana Laha).

## Abstract

This formal comment is in response to “A quest for miRNA bio-marker: a track back approach from gingivo buccal cancer to two different types of precancers” written by De Sarkar and colleagues in 2014. The above-mentioned paper found seven miRNAs to be significantly deregulated in 18 gingivo-buccal cancer samples. However, they suspected more miRNAs to be deregulated based on their exploratory statistical analysis. To control the false discovery rate (FDR), the authors used the Benjamini Hochberg (BH) method, which does not leverage any available biological information on the miRNAs. In this work, we show that some specialized versions of the BH method, which can exploit positional information on the miRNAs, can lead to seven more discoveries with this data. Specifically, we group the closely located miRNAs, and use the group Benjamini Hochberg (GBH) methods (Hu et al., 2010), which reportedly have more statistical power than the BH method (Liu et al., 2019). The whole transcriptome analysis of Sing et al. (2017) and previous literature on the miRNAs suggest that most of the newly discovered miRNAs play a role in oncogenesis. In particular, the newly discovered miRNAs include hsa-miR-1 and hsa-miR-21-5p, whose cancer-related activities are well-established. Our findings indicate that incorporating the GBH method into suitable microarray studies may potentially enhance scientific discoveries via the exploitation of additional biological information.

## Introduction

MicroRNAs (miRNA) are short non-coding RNA molecules, whose oncogenic activities are well known in different types of cancers including oral cancer [1–3]. [4] investigated the functional role of miRNAs in an important subtype of oral squamous cell carcinoma (OSCC) prevalent in South Asia, namely gingivo buccal squamous cell carcinoma (GBSCC) [5]. They studied the miRNA profile of 18 GBSCC patients, and found seven miRNAs to be significantly deregulated in the cancer tissues. Although their list included some previously-reported OSCC-related miRNAs such as hsa-miR-133a, hsa-miR-7, hsa-miR-204, and hsa-miR-31, it excluded many miRNAs that are known to promote tumorigenesis in OSCC. Examples include hsa-miR-1 [6], hsa-miR-181 [7], hsa-miR-99a-3p [8], hsa-miR-17 [9], and hsa-miR-25 [10, 11]. Some of the target genes of these miRNAs are found to be significantly deregulated in [12]’s whole transcriptome analysis, which was based on a cohort with a large overlap with [4]’s cohort. [4]’s exploratory analysis also hinted at the possibility that there were more deregulated miRNAs outside the list of their seven miRNAs. In light of the above, the question that naturally arises is whether [4]’s statistical methods have been overly conservative.

In the first step of their statistical analysis, [4] performed pairwise t-tests to compare the miRNA expressions between cancer affected and the normal (control) tissues. Since there are a total of 522 miRNAs, the latter generates the need for type I error control for simultaneously testing a family of hypotheses. To this end, [4] opted for controlling the false discovery rate (FDR) by using the Benjamini-Hochberg (BH) method [13]. It is well known in the statistical literature that Benjamini-Hochberg (BH) method is conservative, especially when the signal is sparse, i.e., the proportion of null hypotheses is large [14, 15]. Although the above may explain the apparent conservative results in [4], the bigger question for us is this: is it possible to do better than the standard BH method?

The Benjamini-Hochberg (BH) method [13] does not use any prior information on the dependency among the miRNA expressions. However, such information is available to us because closely located miRNAs are more likely to be correlated than distant miRNAs [16, 17]. In particular, miRNAs may form positional clusters depending on their location on the chromosome. There are updated versions of the BH method [14, 18] tailored for such structured and correlated data to yield better statistical power. The goal of this report is to show that such specialized versions of the BH method can lead to the detection of more miRNAs with [4]’s data. The whole transcriptome analysis of [12] and previous literature on the miRNAs suggest that most of the newly discovered miRNAs (we have seven of them) are associated with oncogenesis.

## 1 Materials and methods

### 1.1 Dataset

We consider the dataset used in [4], who made the data publicly available. It contains Δ*Ct* values of 522 miRNA pairs collected from tumor cells and normal (control) cells of 18 GBSCC patients in West Bengal, India. The Δ*Ct* values quantify the miRNA expression levels.

### 1.2 Statistical analysis

We say a miRNA is up/ down-regulated if its expression is *>* 2 fold up/ down regulated. The first step of our statistical analysis is analogous to [4], where they perform one-sided pairwise t-tests at the level of significance 0.05 to test whether miRNA is significantly deregulated. See [4] for more details on these tests. The combined type-I error of the multiple t-tests can be greater than 0.05, which calls for a procedure for type-I error control. To this end, following the common practice of false discovery rate (FDR) control, [4] uses the Benjamini-Hochberg (BH) method, arguably the most popular FDR correction method for microarray data [19–21]. The BH method is provably valid if the tests are independent or weakly dependent [22]. However, it has some limitations.

#### Limitations of BH method

The first criticism of the BH method is that it is overly conservative when the true signals are sparse, i.e., the number of truly deregulated miRNAs is small [14, 15]. We will illustrate this issue with an example. Suppose only ten miRNAs are deregulated in our dataset ([4] found seven deregulated miRNAs) and the miRNA expressions are independent. In this case, it can be shown that if one aims to achieve an FDR of level *α* by applying the BH method, the actual FDR turns out to be approximately 0.019*α* [13]. If we increase the number of deregulated miRNAs in our example to 30 and 50, the true FDR increases to 0.06*α* and *α/*10, respectively, which is still much smaller than the targeted level *α*.

Second, the ordinary BH method can not leverage any prior information on the dependence structure among the hypotheses although such prior information may increase the statistical power [18]. This is an important point for us because we do have prior information on the dependence structure among the hypotheses, leading to a group structure that the BH method [13] can not exploit.

#### Group structure in our data

Since neighboring genes tend to be correlated [16, 17], we group the miRNAs based on their spatial location on the chromosome. We understand that physical proximity may not be the primary reason behind the association among the miRNAs. They may be correlated due to other biological factors, e.g., the transcription factors. However, since our sample size is quite small (only 18 cancer patients), we want to keep the groups as simple as possible. We will see in Section 3 that even this simple grouping strategy leads to the discovery of many relevant miRNAs. Therefore, in this study, we opt for a simple positional grouping, where we place miRNAs in the same group only if their chromosome number, arm, and strand are the same. Since there are 23 chromosomes, each with two strands, and each strand has two arms, a total of 92 such groups are possible. However, we exclude the groups with no membership from our 522 miRNAs, which leaves us with a total of 67 groups. It is also possible, at least in principle, to group the miRNAs using data-dependent clustering methods such as k-means clustering. However, recent statistical literature has begun to highlight that using the same data for clustering and downstream testing can inflate the type I error [23–25]. Also, we do not have any solid criterion for choosing the number of clusters for data-dependent clustering. Therefore, we opt for pre-specified groups instead, which are based on prior biological knowledge.

Although we can create groups in different ways, they are statistically meaningful only when the intra-group association is stronger than the inter-group association. S1 File provides some exploratory analysis, which shows that our data does not refute the above possibility for our groups. We defer further discussion on the groups to S1 file. We will now move on to some FDR control methods that can be fed group-based information to improve the statistical power.

#### Group-based BH method

There is an updated version of the BH method for grouped hypotheses, and this version goes by the name grouped Benjamini Hochberg method (GBH [18]). The GBH method aims to improve the statistical power by recalibrating the BH method by putting more importance on the potentially important groups [14, 18]. The relative importance of a group is measured by the proportion of true null hypotheses in that group. Naturally, a group is more important if the latter proportion is small, and the group Benjamini Hochberg method is more likely to reject hypotheses from this group. When the p-values are independent or weakly correlated, it can be shown that this method yields FDR of approximately level *α* if the group-wise null proportions are estimated reasonably well [14, 18]. We want to make it clear that this method does not use the groups as the unit of rejections. Instead, the rejections are carried out at the level of the individual hypotheses, where the information on the groups is used to improve the power.

The proportion of the null hypotheses in each group can be estimated from the data using different statistical methods, e.g., the two-stage step-up (TST) method of [26], the Least-Slope (LSL) method of [27], and the likelihood-based method of [14]. The respective FDR control methods are known as TST-GBH [18], LSL-GBH [18], and SABHA [14]. The GBH method has previously been applied in many biomedical studies including genetic studies, where the goal was to discover differentially expressed genes in the presence of positional groups [28, 29]. Therefore, alongside BH, we also use TST-GBH, LSL-GBH, and SABHA to control the FDR of the pairwise t-tests. SABHA requires two tuning parameters: (1) *E*, a lower bound on the proportion of true null hypotheses, and (2) *τ*, used for calibrating the individual p-value thresholds. Similiar to [14], we take *E* = 0.1 and *τ* = 0.5. We found that, for our data, the discoveries are not sensitive to small perturbations of either of these tuning parameters.

The GBH methods are believed to [18] have an edge over the BH method when (a) the majority of hypotheses are null, i.e., most miRNAs are not deregulated in cancer tissues (b) the groups are heterogeneous in that the proportion of nulls are different across the groups. [4]’s findings indicate that (a) probably holds for our data, whereas (b) is harder to verify. However, since our groups are physically separated, and, as mentioned earlier, the within-group association is estimated to be stronger than the between-group association (see File S1 for the analysis), it is unlikely that our groups are homogeneous. We remark that there are many other FDR control methods tailored for dependent hypotheses [15, 30, 31], but methods targeting broad classes of dependence structures generally fail to leverage information on specific structures such as the group structure fully [14, 32].

### Missing observations

Some miRNAs did not express in all 18 patients, but interestingly the missingness always occurs in pairs. To elaborate, these miRNAs are expressed in neither the cancer tissues nor the control tissue in those subjects. Figure S2 displays the histogram of the missing pairs for our miRNAs. Close inspection shows that 69% of the miRNAs have at least one pair of missing observations. The missingness may occur from low expression of the miRNAs in some individuals, but it is hard to pinpoint the exact reason since some information was also lost during the pruning of the miRNAs [4]. Anyway, it is clear that the missingness is not at random. [4] replaced the missing observations using median imputation; see [4] for more details. However, imputing the missing values with mean or median can artificially decrease the sample variance in small samples, which can lead to false positives [33]. Since the missing values always occur in pairs in our data, median imputation leads to the inclusion of identical pairs, which leaves us at a higher risk of artificial variance deflation.

We found that the median imputation leads to a sharp increase in the number of discoveries irrespective of the FDR control method. Table S3 lists the miRNAs detected using the imputed data. Alarmingly, many of the extra discoveries correspond to miRNAs with moderate to severe missingness (cf. Table S3), hinting that these discoveries may be the result of the imputation-driven variance deflation. The picture is similar for mean imputation. Since imputation yields a strikingly different result, we decide to avoid it altogether in our main analysis. Doing so does not cause any loss of information because the missingness always occurs in pairs.

## 2 Results

Table 1 displays the deregulated miRNAs detected by the TST-GBH, LSL-GBH, SABHA, and the BH method. They reject 15, 5, 5, and 3 hypotheses, respectively. Table 1 indicates that TST-GBH is more liberal than other GBH methods, which was previously noted by [34]. The other two GBH methods, LSL-GBH and SABHA, mostly agree with each other. Table 1 contains the original seven miRNAs detected by [4], and seven more new miRNAs. Among the newly detected miRNAs, hsa-miR-1247-5p, hsa-miR-1, hsa-miR-99a-3p, and hsa-miR-486 3p/ 5p are downregulated, and hsa-miR-21-5p and hsa-miR-147b are upregulated in the cancer tissues. A heatmap for the ΔΔ*Ct* values of these miRNAs is also constructed (see Figure 1). Although these new miRNAs were not technically detected by [4], most of them were mentioned in [4]. [4]’s cluster analysis found a cluster of 13 patients, who had 30 miRNAs deregulated in the cancer tissues (see Table 6 of [4]). Except for hsa-miR-486-3p/5p, all other newly detected miRNAs belong to this set of 30 miRNAs. In particular, the newly detected miRNAs include hsa-miR-1, which [4] speculated to be deregulated, and hsa-miR-21-5p, which is an established oncogene. We will discuss more on the newly detected miRNAs in Section 3. The R code for reproducing the statistical analysis in this paper can be found at https://github.com/SalilKoner/miRNA-biomarker.

**Table 1.** Significantly deregulated miRNAs.

| | MiRNA | Chr. | strand | $\Delta\Delta Ct$ | p-value | Missing<br>pairs | Significant tests |
| --- | --- | --- | --- | --- | --- | --- | --- |
| 1 | hsa-miR-133a-3p* | chr18 | - | 6.7 | 0.00 (↓) | 0 | BH, TST, LSL, SABHA |
| 2 | hsa-miR-206* | chr6 | + | 6.0 | 0.00 (↓) | 0 | BH, TST, SABHA |
| 3 | hsa-miR-31-3p* | chr9 | - | -3.8 | 0.00 (↑) | 2 | BH, TST, LSL, SABHA |
| 4 | hsa-miR-1293* | chr12 | - | -4.8 | 0.00 (↑) | 9 | TST |
| 5 | hsa-miR-1247-5p | chr14 | - | 2.4 | 0.01 (↓) | 2 | TST |
| 6 | hsa-miR-147b | chr15 | + | -2.1 | 0.01 (↑) | 2 | TST |
| 7 | hsa-miR-7-5p* | chr15 | + | -3.1 | 0.00 (↑) | 2 | TST, LSL |
| 8 | hsa-miR-21-5p | chr17 | + | -2.2 | 0.00 (↑) | 0 | TST |
| 9 | hsa-miR-1 | chr18 | - | 5.2 | 0.00 (↓) | 0 | TST, LSL, SABHA |
| 10 | hsa-miR-99a-3p | chr21 | + | 2.8 | 0.01 (↓) | 0 | TST |
| 11 | hsa-miR-486-3p | chr8 | - | 2.4 | 0.00 (↓) | 0 | TST |
| 12 | hsa-miR-486-5p<sup>†</sup> | chr8 | - | 2.0 | 0.02 (↓) | 0 | TST |
| 13 | hsa-miR-31-5p* | chr9 | - | -3.4 | 0.00 (↑) | 0 | TST, LSL, SABHA |
| 14 | hsa-miR-204-5p* | chr9 | - | 4.6 | 0.00 (↓) | 2 | TST |

These miRNAs were detected by at least one of the methods among BH, TST-GBH, LSL-GBH, and SABHA. In this table, we abbreviate TST-GBH and LSL-GBH by TST and GBH, respectively. The columns “Chr.” and “strand” indicate which chromosome and strand the miRNA belongs to, respectively. The column $\Delta\Delta Ct$ gives the $\Delta\Delta Ct$ values for each miRNA. We report the raw p-values before the FDR correction because TST-GBH, LSL-GBH, and SABHA do not yield FDR-corrected p-values. The signs after the p-values indicate whether the miRNAs were upregulated (↑), or downregulated (↓). The column “Missing pairs” gives the number of missing pairs of observations for each miRNA. The column “Significant tests” indicates which methods detected the miRNA to be significantly deregulated.
\* These genes were reported to be deregulated in [4].

**Fig 1.**
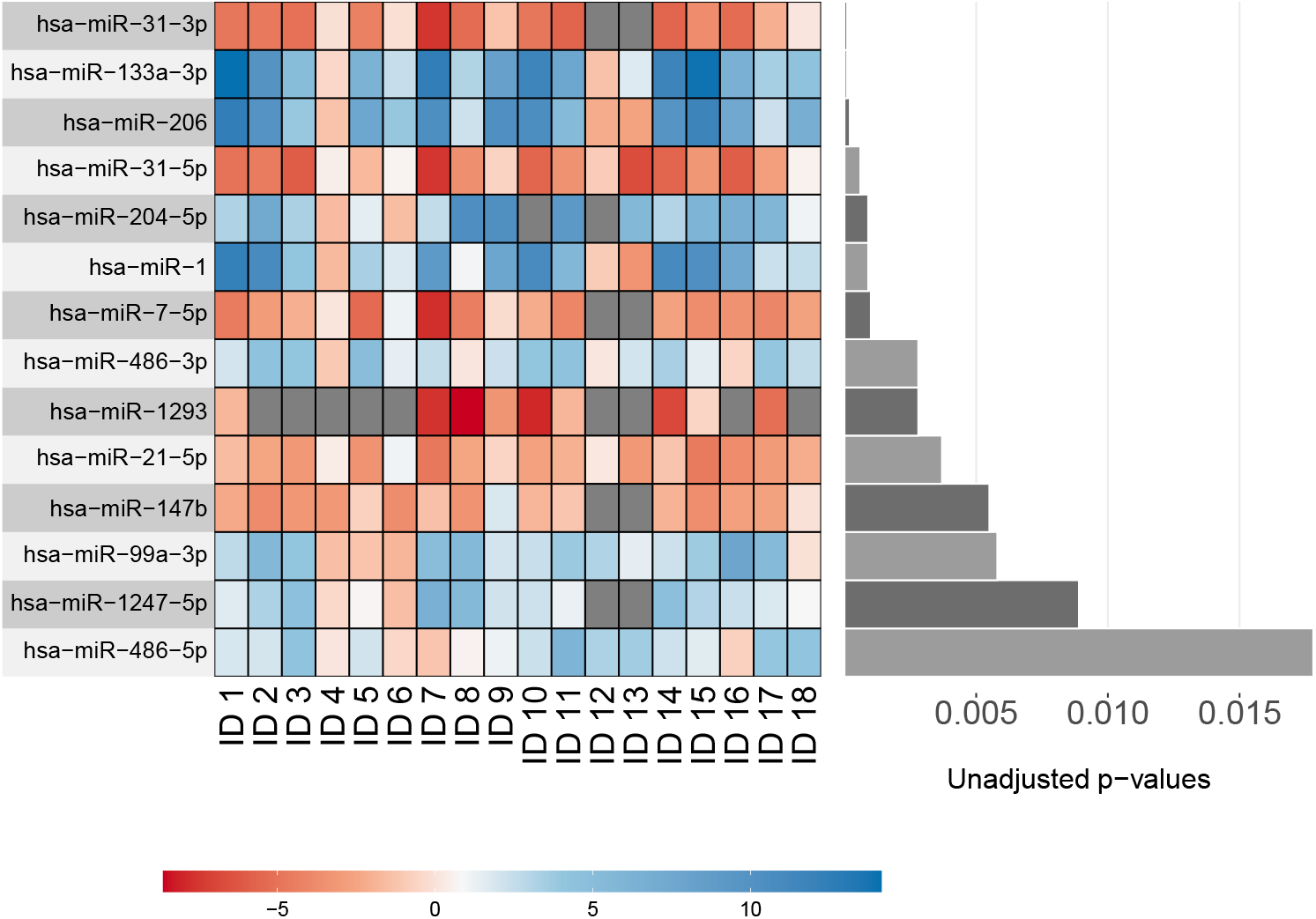
Heatmap of the ΔΔ*CT* values. The heatmap is based on the 18 patients and all discovered miRNAs from Table 1. The miRNAs are placed in the ascending order of the raw/ unadjusted p-values obtained from the paired t-tests.

Table 1 also displays the missingness pattern of the discovered genes. Except for hsa-miR-1293, the other deregulated miRNAs have at most two missing pairs, which amounts to only around 11% of the data. The interesting case is hsa-miR-1293 – 50% of the data was missing for this miRNA. Although this amount of missingness raises doubts regarding any conclusion on this miRNA, hsa-miR-1293 is known to promote cell migration and invasion in renal cell carcinoma [35]. See [4] for more discussion on this miRNA.

## 3 Discussion

As we will see, the newly discovered miRNAs show a similar direction of expression deregulation as found in previous literature. Moreover, some target genes of these newly discovered miRNAs are deregulated in the opposite direction compared to their targeting miRNAs in [12]’s whole transcriptome analysis (see Table 2). We remind the readers that the latter is based on a cohort of twelve cancer patients that included ten of [4]’s subjects. Below we discuss the newly discovered miRNAs in further detail.

**Table 2.** The target genes of the newly discovered miRNAs that are deregulated in [12]’s study.

| miRNA | deregulated target genes found by [12] | references |
| --- | --- | --- |
| hsa-miR-21-5p | MEF2C (↓), PDCD4 (↓), TIMP3 (↓), PPARA (↓)<br>RECK (↓), SPRY1 (↓), SPRY2 (↓), THRB (↓) | [36] |
| hsa-miR-1 | FAM101B (↑), TRANK1 (↑)<br>ZNF281 (↑), Runx2 (↑) CXCR4 (↑) | [37–40] |
| hsa-miR-486-3p | FLNA (↑) | [41] |
| hsa-miR-486-5p | KIAA1199 (↑) | [41, 42] |
| hsa-miR-99a-3p | BCAT1 (↑), MTHFD2 (↑), RRM2 (↑) | [8, 43] |

The arrows next to the genes indicate the direction of the deregulation. The information on the target genes for each miRNA was obtained from the corresponding references.

### hsa-miR-21-5p, upregulated

hsa-miR-21 is claimed to be a ubiquitous oncogene [44]. Although its cancer-promoting activities in OSCC have been verified, most studies in this context predominantly consider tongue or floor-of-the-mouth cancer samples [10, 11]. Eight target genes of hsa-miR-21 [36] are found to be downregulated in [12], which include MEF2C, PDCD4, TIMP3, PPARA, RECK, SPRY1, SPRY2, and THRB. Notably, PDCD4 is one of the main target genes of hsa-miR-21 [45]. hsa-miR-21 reportedly suppresses the activities of these genes [36]. Downregulation of these genes is linked to different hallmarks of cancer [46], such as resisting cell death (TIMP3, PDCD4, PPARA, TIMP3, SPRY 1/2), activating invasion and metastasis (PPARA, TIMP3, SPRY 1/2, RECK), tumor-promoting inflammation (PDCD4), and inducing angiogenesis (TIMP3, RECK) [36]. hsa-miR-21 has been advocated by many to be a prognostic biomarker for oral squamous cell carcinoma (OSCC) [1, 36, 47, 48]. Our findings do not refute this claim at least for GBSCC.

### hsa-miR-1, downregulated

hsa-miR-1 plays a critical role in physiological processes in the smooth and skeletal muscles [38]. Numerous studies verified its role as a tumor suppressor [39], whose downregulation can potentially cause several types of cancers including OSCC [6]. Target oncogenes of hsa-miR-1 that are upregulated in [12] include RUNX2, ZNF281, and CXCR4. Since the expression of these genes is modulated by hsa-miR-1 [39, 40, 49, 50], we speculate that their upregulation is the consequence of the downregulation of hsa-miR-1 in the cancer cells. Oncogenetic activities of these genes are well-established in the literature, e.g., RUNX2 in breast cancer [51, 52], CXCR4 in thyroid cancer, and osteosarcoma [53], and ZNF281 in pancreatic Cancer [54].

### hsa-miR-486-3p and hsa-miR-486-5p, downregulated

Various studies have established hsa-miR-486 to be a tumor suppressor [55]. In many cancers including OSCC, its downregulation is associated with migration and invasion [41]. FLNA and KIAA1199, which are target genes of hsa-miR-486-3p and hsa-miR-486-5p, respectively, were upregulated in [12], which is unsurprising since the respective miRNAs suppress these genes [42, 56]. FLNA promotes cell migration in Laryngeal squamous cell carcinoma [41] where KIAA1199 is a well-known promoter of invasion and metastasis in thyroid cancer [42].

### hsa-miR-99a-3p, downregulated

Studies have suggested that hsa-miR-99a-3p may act as a tumor suppressor in some types of carcinoma [43]. Among the target oncogenes of hsa-miR-99a-3p, BCAT1, MTHFD2, and RRM2 are upregulated in [12]. Studies found these oncogenes to be associated with poor cancer prognosis [8, 43]. hsa-miR-99a-3p has been found to suppress the oncogenes BCAT1 and MTHFD2 in head and neck squamous cell carcinoma [8], and the oncogene RRM2 (direct target) in renal carcinoma [43].

### hsa-miR-1247-5p, downregulated and hsa-miR-147b, upregulated

hsa-miR-1247 acts as a tumor suppressor and is reported to be downregulated in several cancers [57–60]. In contrast, hsa-miR-147b promotes cell proliferation and activates invasion [61], and is found to be upregulated in several types of lung cancers [62–65]. To the best of our knowledge, well-known target genes of hsa-miR-1247-5p and hsa-miR-147b did not exhibit deregulation in [12]. However, RCC1, which is structurally similar to a target gene of hsa-miR-1247-5p, RCC2 [66], is upregulated in [12]. hsa-miR-1247-5p is known to suppress the oncogenic activities of RCC2, which is found to promote tumor prognosis in pancreatic cancer [58]. RCC1 is also a known promoter of tumor progression, however [67].

### Overlap with prior studies

To the best of our knowledge, among our discovered miRNAs, hsa-miR-7-5p, hsa-miR-204-5p, hsa-miR-21, and miR-99a-3p have the most notable mention in prior studies in OSCC [1–3]. Note that hsa-miR-21 and miR-99a-3p belong to the new discoveries that have been made possible by the GBH method. However, our methods still failed to detect many miRNAs that were found to be significantly deregulated in some of the above-mentioned studies. The lack of overlap may be due to the limited sample size of our data. Many statistical methods have such issues with multicollinearity [68]. However, most of these studies [1–3] mainly focus on tongue and floor-of-the-mouth cancer patients, whose clinical behaviors as well as molecular and immunoproteome profile may be different from GBSCC, a cancer predominant in the tobacco-chewing population of South Asia [5]. This can also be a potential reason behind the absence of many previously reported miRNAs from our list.

## 4 Conclusion

The grouped BH methods are able to detect some of the miRNAs that were (a) undetected but speculated to be deregulated in [4] or (b) previously known to play a role in OSCC. The new discoveries may be attributable to the gain in the statistical power obtained by the incorporation of positional information on the miRNAs. Similar to [28, 29], our results suggest that using specialized BH methods in genetic data with pre-existing group structures may help the cultivation of novel information that would otherwise remain hidden. However, future experiments are required to shed further light on this speculation.

## 5 Acknowledgments

The authors are grateful to Michele Shaffer for her helpful comments and advice.

## Supporting information

**S1 File. Details on the groups**. Contains (1) details on the size and the membership of the groups and (2) an exploratory analysis to compare the intra-group and the inter-group correlations. (PDF)

**S2 Fig. Histogram of the number of missing pairs of miRNAs**. The x-axis corresponds to the number of missing pairs, and the y-axis corresponds to the frequency of the miRNAs with missing pairs. (PDF)

**S3 Table. Significantly deregulated miRNAs with the median-imputed data**. (PDF)

R codes to reproduce all the tables and the figures presented in the manuscript as well as in the supporting information document are deposited at https://github.com/SalilKoner/miRNA-biomarker

