## Supplement for "Discovery of new deregulated miRNAs in gingivo buccal carcinoma using Group Benjamini Hochberg method: a commentary on “A quest for miRNA bio-marker: a track back approach from gingivo buccal cancer to two different types of precancers”"

### S1 File: Details on the groups

Table S1 tabulates the frequency (number of members) as well as the physical size of each group. The size is calculated by taking the difference between the start coordinates of the most extremely situated chromosomes in each group. Note that taking the above difference makes sense because the miRNAs in a group are situated on the same arm of the same chromosome. Table S1 reveals that most physically large groups have small to moderate frequency. Therefore, albeit their large size, further splits of these groups seem to be unnecessary. On the other hand, there are some thinly populated groups. The number of such groups can be decreased by merging the groups on the two arms of the same strand. However, doing so leads to a nominal difference – only hsa-miR-486-5p ceases to be statistically significant after the merging. Another interesting observation from Table 1 and Table S1 is that membership in overly populated groups does not increase the chances of being detected. In fact, all discovered miRNAs belong to groups with frequencies less than ten.

**Table S1.** lists the size, frequency, chromosome number, and the corresponding strand for each group. The physical size of each group is calculated by taking the difference between the start co-ordinates (scaled by $10^5$) of the most extremely located microRNAs in the groups. Here frequency corresponds to the number of miRNAs in a group. There can be at most two groups on the same strand of the same chromosome because there are only two arms, each of which corresponds to a different group.

| Group | Size | Frequency | Chr. | Strand |
|---|---|---|---|---|
| 1 | 89.30 | 9 | chr1 | - |
| 2 | 66.21 | 19 | chr1 | - |
| 3 | 116.53 | 10 | chr1 | + |
| 4 | 58.09 | 3 | chr1 | + |
| 5 | 0.06 | 2 | chr10 | - |
| 6 | 47.04 | 4 | chr10 | - |
| 7 | 5.80 | 2 | chr10 | + |
| 8 | 59.69 | 4 | chr10 | + |
| 9 | 1.59 | 4 | chr11 | - |
| 10 | 57.36 | 11 | chr11 | - |
| 11 | 67.78 | 6 | chr11 | + |
| 12 | 71.94 | 8 | chr12 | - |
| 13 | 6.00 | 5 | chr12 | + |
| 14 | 43.23 | 8 | chr12 | + |
| 15 | 57.56 | 5 | chr13 | - |
| 16 | 1.12 | 13 | chr13 | + |
| 17 | 78.17 | 4 | chr14 | - |
| 18 | 38.65 | 55 | chr14 | + |
| 19 | 54.96 | 9 | chr15 | - |
| 20 | 43.43 | 7 | chr15 | + |
| 21 | 65.10 | 2 | chr16 | - |
| 22 | 14.92 | 5 | chr16 | + |
| 23 | 13.07 | 3 | chr16 | + |
| 24 | 17.63 | 13 | chr17 | - |
| 25 | 51.91 | 17 | chr17 | - |

Continued on next page

**Table S1 – continued from previous page**

| Group | Size | Frequency | Chr. | Strand |
|---|---|---|---|---|
| 26 | $5.7 \times 10^{-5}$ | 2 | chr17 | + |
| 27 | 29.47 | 5 | chr17 | + |
| 28 | 14.08 | 3 | chr18 | - |
| 29 | 18.28 | 8 | chr19 | - |
| 30 | 3.86 | 3 | chr19 | - |
| 31 | 3.98 | 5 | chr19 | + |
| 32 | 8.31 | 37 | chr19 | + |
| 33 | 14.27 | 3 | chr2 | - |
| 34 | 87.14 | 3 | chr2 | - |
| 35 | 104.97 | 8 | chr2 | + |
| 36 | 23.35 | 3 | chr20 | - |
| 37 | 58.65 | 7 | chr20 | + |
| 38 | 9.03 | 6 | chr21 | + |
| 39 | 24.21 | 4 | chr22 | - |
| 40 | 26.49 | 10 | chr22 | + |
| 41 | 41.87 | 8 | chr3 | - |
| 42 | 77.29 | 2 | chr3 | - |
| 43 | 12.20 | 4 | chr3 | + |
| 44 | 83.59 | 10 | chr3 | + |
| 45 | 22.53 | 3 | chr4 | - |
| 46 | 29.90 | 6 | chr4 | - |
| 47 | 38.53 | 4 | chr4 | + |
| 48 | 115.95 | 5 | chr4 | + |
| 49 | 3.75 | 2 | chr5 | - |
| 50 | 126.19 | 9 | chr5 | - |
| 51 | 26.99 | 7 | chr5 | + |
| 52 | 85.42 | 4 | chr6 | - |
| 53 | 0.01 | 2 | chr6 | + |
| 54 | 26.15 | 6 | chr7 | - |
| 55 | 37.45 | 17 | chr7 | - |
| 56 | 8.82 | 2 | chr7 | + |
| 57 | 77.33 | 7 | chr7 | + |
| 58 | 30.63 | 5 | chr8 | - |
| 59 | 45.07 | 8 | chr8 | - |
| 60 | 127.40 | 3 | chr8 | + |
| 61 | 7.38 | 3 | chr9 | - |
| 62 | 57.58 | 4 | chr9 | - |
| 63 | 118.85 | 14 | chr9 | + |
| 64 | 7.98 | 6 | chrX | - |
| 65 | 77.69 | 32 | chrX | - |
| 66 | 41.68 | 9 | chrX | + |
| 67 | 70.39 | 5 | chrX | + |

#### Exploratory analysis to compare the inter-group and intra-group correlation

The goal of this section is to compare the intra-group and the inter-group correlations in our data. Vaguely speaking, these two terms measure the association within and across the groups, respectively. However, to rigorously compare these correlations, we need a formal way to define these quantities. To rigorize the setup, we posit a simple random effect model on the $\Delta\Delta CT$ values [1], which helps us to quantify and estimate the inter-group and intra-group correlations. Using a random effect model to estimate the intra-class and inter-class correlations is not uncommon in statistic literature [2]. It should be kept in mind that this model can be misspecified. Even then, the estimators of the inter-group and the intra-group correlations may still provide realistic assessments of the strength of the corresponding associations.

To formalize the setup, we will introduce some new notations at first. Denote by $\{Y_{ij} : j = 1, \ldots, 522\}$ the $\Delta\Delta CT$ values of the $j$th miRNA of the $i$th subject/ patient. Let $g(j)$ be the group indicator of the $j$th miRNA, e.g., if the $j$th miRNA belongs to the first group, then $g(j) = 1$. The group index $g(j)$ takes value between 1 and 67.

We will posit a random effect model [3] on the $\Delta\Delta CT$ values as

$$Y_{ij} = \mu + S_i + G_{ig(j)} + E_{ij},$$

where $\mu$ is the overall mean, $S_i$ is the random effect of the ith patient, $G_{ig(j)}$ is the random effect of the $g(j)$th group for the $i$th patient, and $E_{ij}$ is the random error for the $i$th subject and the $j$th miRNA. The random error can represent independent measurement errors or other underlying biological factors. We let the collections of random variables $\{S_i : i = 1, \ldots, 18\}$, $\{G_{ig(j)} : 1 \leq i \leq 18,\ 1 \leq j \leq 522\}$, and $\{E_{ij} : 1 \leq i, j \leq 522\}$ to be independent of each other. We will also assume that $S_i \overset{\text{iid}}{\sim} N(0, \sigma_s^2)$, $G_{ig(j)} \overset{\text{iid}}{\sim} N(0, \sigma_g^2)$, and $E_{ij} \overset{\text{iid}}{\sim} N(0, \sigma_e^2)$.

Under the above model, the correlation between the $\Delta\Delta CT$ values of two miRNAs belonging to two different groups within the same subject is unique. The above correlation will thus be a natural candidate for the inter-group correlation. Straight-forward algebra shows that the inter-group correlation equals $\sigma_s^2/(\sigma_s^2 + \sigma_g^2 + \sigma_e^2)$. Likewise, under our model, the $\Delta\Delta CT$ values of two miRNAs belonging to the same group within the same subject have identical correlations across the subjects and the miRNAs – this correlation will be a natural candidate for the intra-group correlation. The intra-group correlation is given by $(\sigma_s^2 + \sigma_g^2)/(\sigma_s^2 + \sigma_g^2 + \sigma_e^2)$.

We fit the simple random-effect model using SAS proc mixed [4] to obtain an estimate of the inter-group correlation as 0.04, which suggests a weak association between the $\Delta\Delta CT$ values between groups. The magnitude of the intra-group correlation estimated from the above model is 0.14, which is noticeably larger than the inter-group correlation. Therefore, our model suggests that the inter-group correlation is perhaps weaker than the intra-group correlation.

4. SAS Institute Inc. SAS/STAT® 9.2 User's Guide. Cary, NC: SAS Institute Inc; 2008.

### S2 Fig.

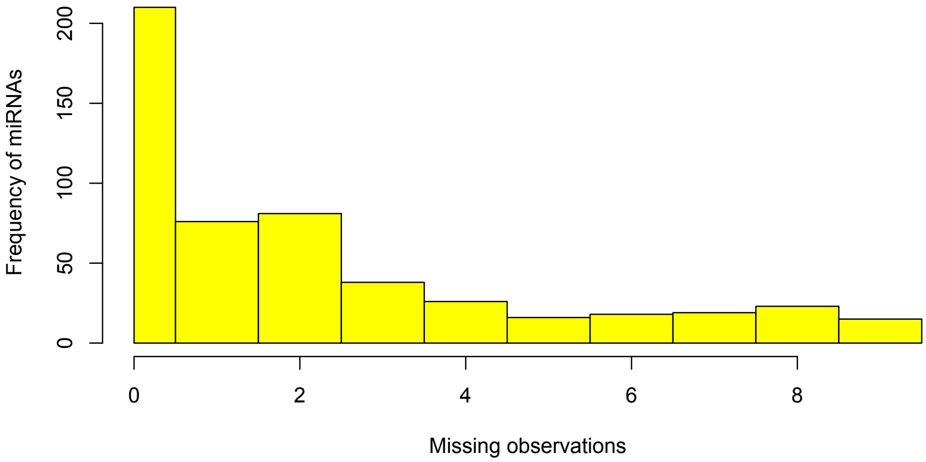


**Fig 1. Histogram of the number of missing pairs of miRNAs**. Here we considered the 522 miRNAs from our analysis. The x-axis corresponds to the number of missing pairs, and the y-axis corresponds to the frequency of miRNAs with missing pairs.

### S3 Table: Median imputation

**Table S2.** Significantly deregulated miRNAs with the median-imputed data

| MiRNA | chr. | strand | $\Delta\Delta$CT | p-value | Missing pairs | Significant tests |
|---|---|---|---|---|---|---|
| hsa-miR-548k* | chr11 | + | -2.7 | 0.00 ($\uparrow$) | 8 | BH, TST, SABHA |
| hsa-miR-1293 | chr12 | - | -4.8 | 0.00 ($\uparrow$) | 9 | BH, TST, SABHA |
| hsa-miR-7-5p | chr15 | + | -3.1 | 0.00 ($\uparrow$) | 2 | all tests |
| hsa-miR-133a-3p | chr18 | - | 6.7 | 0.00 ($\downarrow$) | 0 | all tests |
| hsa-miR-1 | chr18 | - | 5.2 | 0.00 ($\downarrow$) | 0 | all tests |
| hsa-miR-206 | chr6 | + | 6.0 | 0.00 ($\downarrow$) | 0 | BH, TST, SABHA |
| hsa-miR-133b* | chr6 | + | 2.7 | 0.00 ($\downarrow$) | 8 | BH, TST, SABHA |
| hsa-miR-31-3p | chr9 | - | -3.8 | 0.00 ($\uparrow$) | 2 | all tests |
| hsa-miR-31-5p | chr9 | - | 3.4 | 0.00 ($\uparrow$) | 0 | all tests |
| hsa-miR-204-5p | chr9 | - | 4.6 | 0.00 ($\downarrow$) | 2 | BH, TST, SABHA |
| hsa-miR-504-5p* | chrX | - | 4.7 | 0.00 ($\downarrow$) | 6 | all tests |
| hsa-miR-891a-5p* | chrX | - | 4.4 | 0.00 ($\downarrow$) | 7 | all tests |
| hsa-miR-508-3p* | chrX | - | 4.0 | 0.00 ($\downarrow$) | 6 | all tests |
| hsa-miR-1290* | chr1 | - | -2.2 | 0.00 ($\uparrow$) | 2 | TST |
| hsa-miR-135b-3p* | chr1 | - | -2.2 | 0.00 ($\uparrow$) | 2 | TST |
| hsa-miR-200b-5p* | chr1 | + | -3.5 | 0.00 ($\uparrow$) | 3 | TST |
| hsa-miR-200c-5p* | chr12 | + | -2.3 | 0.01 ($\uparrow$) | 3 | TST |
| hsa-miR-1247-5p | chr14 | - | 2.4 | 0.00 ($\downarrow$) | 2 | TST, LSL |
| hsa-miR-211-5p* | chr15 | - | 3.7 | 0.00 ($\downarrow$) | 4 | TST, LSL |
| hsa-miR-147b | chr15 | + | -2.1 | 0.00 ($\uparrow$) | 2 | TST, LSL |
| hsa-miR-21-5p | chr17 | + | -2.2 | 0.00 ($\uparrow$) | 0 | TST |
| hsa-miR-99a-3p | chr21 | + | 2.8 | 0.01 ($\downarrow$) | 0 | TST, LSL |
| hsa-miR-383-5p* | chr8 | - | 2.7 | 0.03 ($\downarrow$) | 4 | TST |
| hsa-miR-486-3p | chr8 | - | 2.4 | 0.00 ($\downarrow$) | 0 | TST, LSL |
| hsa-miR-486-5p | chr8 | - | 2.0 | 0.02 ($\downarrow$) | 0 | TST |
| hsa-miR-770-5p* | chr14 | + | 3.0 | 0.00 ($\downarrow$) | 3 | LSL |
| hsa-miR-299-5p* | chr14 | + | 2.5 | 0.00 ($\downarrow$) | 4 | LSL |

Table of the significantly deregulated miRNAs detected using the median imputed data. These miRNAs were detected by at least one of the four methods: BH, SABHA, TST-GBH, and LSL-GBH. (The last two methods are shown in the table as TST and LSL, respectively.) The columns "Chr." and "strand" indicate which chromosome and strand the miRNA belongs to, respectively. The column $\Delta\Delta Ct$ gives the $\Delta\Delta Ct$ values for each miRNA. We report the raw p-values before the FDR correction because the GBH methods, e.g., TST-GBH, LSL-GBH, and SABHA, do not yield FDR-corrected p-values. The signs after the p-values indicate whether the miRNAs were upregulated ($\uparrow$), or downregulated ($\downarrow$). The column "Missing pairs" gives the number of missing pairs of observations for each miRNA. The column "Significant tests" indicates which methods detected the miRNA to be significantly deregulated.
* These miRNAs did not appear in Table 1, which is based on the non-imputed data. We have thirteen such miRNAs in total. Note that these miRNAs also have a relatively higher amount of missingness.
